# Megakaryopoiesis in Dengue virus infected K562 cell promotes viral replication which inhibits endomitosis and accumulation of ROS associated with differentiation

**DOI:** 10.1101/2020.06.25.172544

**Authors:** Jaskaran Kaur, Yogita Rawat, Vikas Sood, Deepak K. Rathore, Shrikant K. Kumar, Niraj K. Kumar, Sankar Bhattacharyya

**Affiliations:** Translational Health Science and Technology Institute, NCR Biotech Science Cluster, PO Box# 4, Faridabad-Gurgaon expressway, Faridabad, Haryana-121001, India; Department of Biochemistry, School of Chemical and Life Sciences, Jamia Hamdard (Hamdard University) Hamdard Nagar, New Delhi - 110062, India

**Keywords:** Dengue virus replication, Megakaryopoiesis, Reactive oxygen species, Endomitosis

## Abstract

In the human host blood Monocytes and bone marrow Megakaryocytes are implicated as major sites supporting high replication. The human K562 cell line supports DENV replication and represent Megakaryocyte-Erythrocyte progenitors (MEP), replicating features of *in vivo* Megakaryopoiesis upon stimulation with Phorbol esters. In this article, we report results that indicate the mutual influence of Megakaryopoiesis and DENV replication on each other, through comparison of PMA-induced differentiation of either mock-infected or DENV-infected K562 cells. We present data showing PMA-induced differentiation to drastically increase DENV replication and a concomitant augmented secretion of infectious virus. Although the mechanism is not clear yet, we show that it is not through an increased uptake of virus by differentiated cells. On the other hand, DENV replication in cells undergoing PMA-induced differentiation, interferes with major differentiation markers of Megakaryopoiesis including activation of ERK1/2 MAP Kinase, endomitosis and surface expression of platelet-specific proteins without any drastic effect on cell death. Among signaling intermediaries of the JAK-STAT pathway, we observed infection associated degradation of SOC3 protein similar to earlier observations with STAT2. DENV infection leads to accumulation of Reactive-oxygen species (ROS) in different cells including K562. PMA-induced differentiation of uninfected K562 cells also leads to intracellular ROS accumulation. Interestingly, we observed ROS accumulation to be suppressed by concomitant DENV replication in K562 cells undergoing PMA-induced differentiation. This is the first report of a model system where DENV replication suppresses intracellular ROS accumulation. The implications of these results for Megakaryopoiesis and viral replication would be discussed.

## Introduction

Platelets play a unique role in tissue homeostasis and regulation of inflammation. The abundance of these small, anuclear cells produced from specialized cells called Megakaryocytes (MKs) in the bone marrow (BM) is controlled by a steady rate of biogenesis (about 1×10^9^/day) and decay, with an average life of 7 days in humans (1). A perturbation of platelet homeostasis is associated with infection by multiple human pathogenic viruses (2). Infection by Dengue virus (DENV) a member of the Flaviviridae, can cause an acute febrile illness with the potential to turn fatal. Beside humans, DENV can infect mosquitoes and these transmit infectious virus along with salivary fluid during a blood meal. Only about a quarter of infected humans exhibit clinical symptoms that include high fever, retro-orbital pain, muscle pain, thrombocytopenia or drop in platelet level etc. that subside after 4-7 days. Subsequently a proportion of symptomatic individuals develop symptoms of severe Dengue characterized by leakage of fluid from the blood vessels, leading to reduction in blood volume in addition to acute thrombocytopenia (3). Multiple mechanisms working in parallel have been suggested to contribute to the thrombocytopenia associated with Dengue infection which include inhibition of platelet biogenesis by infection of Megakaryocytes, augmented platelet decay through either a direct interaction with the virus or upon binding of anti-platelet antibodies (4).

Depending on the cell type the entry of DENV can be mediated by different surface receptors and the complex is endocytosed into endosome following virus binding (5) Acidification of these endosomes by their membrane resident proton pumps induce a fusion between the viral and endosomal membranes leading to release of the viral genome into host cytosol. The genome of DENV is a ∼11 kb long single-stranded plus-sense RNA coding for one large polyprotein which is then cleaved to generate the structural (Core or C, Envelop or E and precursor-Membrane or prM) and non-structural proteins (NS1, NS2A, NS2B, NS3, NS4A, NS4B and NS5) of the virus. After few rounds of translation the viral genomic RNA undergoes replication in **r**eplication **c**omplexes (RC) that are associated with the host cell **E**ndoplasmic **r**eticulum (ER). The NS5 protein performs crucial function in replication of the genome by polymerizing both negative and positive strand genomic RNA through asymmetric replication in favor of the later. Among these only the plus-sense genomic RNA is capped by a Methyl-transferase (MTase) domain also harbored in the NS5, followed by encapsidation of the genomic RNA by the Core or C protein. The newly assembled virus particles contain a RNA-protein complex consisting of a single copy of the genomic RNA with multiple copies of the C protein, encased in a membrane studded with E and prM protein. These transit through the protein secretory pathway of the host cell where the prM protein is cleaved by a host protease called Furin to produce M protein. The virion particles containing M protein along with E in the envelope are released from the host cell without lysis (6).

DENV can infect multiple cell types in the human body including monocytes in the blood, megakaryocytes in the BM and hepatocytes in the liver (5, 7). In addition to these DENV can activate platelets, through a direct interaction with viral receptors on the cell surface (5, 8, 9). Platelets activated by DENV secrete a number of cytokines and chemokines among which CXCL4 promotes viral replication in infected Monocytes, upon binding to CXCR3 receptors on these cells (10). Myelosuppression or a reduction in the mass of BM cells is one of the hallmarks of DENV infection and cells of Megakaryocytic cells have been shown to be particularly permissive to viral replication (11-14). This suggested MK cells to significantly contribute to viremia beside monocytes. In addition the survival of megakaryocytes have been suggested to be negatively affected by infection, thereby leading to reduced platelet biogenesis which contributes to DENV associated thrombocytopenia (15).

Hematopoietic stem cells (HSC) in the BM differentiate into multiple lineages producing all cell types in the blood. One of such lineages are composed of bi-potential Megakaryocyte-Erythrocyte progenitor (MEP) cells that can differentiate into either cell type (16, 17). Differentiation of MEP into a MK involves profound cellular and molecular changes including cell expansion, endomitosis which generates a multi-lobed and polyploid nucleus and expression of platelet specific surface markers on the plasma membrane which will become part of the platelets membrane (18, 19). Platelets bud off from the MK along with a bit of its cytoplasm, containing granules of different types and a host of specific mRNAs (20). Differentiation of MEPs into MKs is promoted by a number of cytokines, the most characterized among which is Thrombopoietin or TPO that binds to the TPO-receptor c-Mpl on the cell surface (21). TPO binding to c-Mpl activates Janus Kinase 2 (JAK2) receptor tyrosine kinase which in turn transduces the signal through specific **S**ignal **t**ransducer and **a**ctivator of **t**ranscription or STATs (STA3 and STAT5), **M**itogen-**a**ctivated **p**rotein **k**inases or MAPK (ERK1/2) and the Phosphatidyl Inosine-3 or PI-3 Kinase (22). STAT3 and STAT5 have been shown to have opposing function in Megakaryopoiesis with the former promoting it (23, 24). Activation of JAK2 pathway increases expression of pro-inflammatory genes and in fact activating these genes through exogenous activation of Toll-like receptor 2 or TLR2 has been shown to promote megakaryocyte development (25). In a similar manner, the activation of ERK1/2 MAPK and inhibition of p38-MAPK promotes Megakaryopoiesis by playing a crucial role in endomitosis (26). Transduction of the signal from JAK2 leads to activated transcription by a number of transcription factors (TFs) increasing the expression of specific genes (27, 28). TFs like GATA-1 and RUNX1 play crucial role in endomitosis through upregulation of CyclinD1 (29, 30). In addition to these different specific pathways like ER-stress and Autophagy have been shown to play role in Megakaryopoiesis (31). A controlled form of apoptosis is known to be crucial for production of platelets from megakaryocytes and ER-stress has been suggested to play a role in this apoptosis (31-33). In the BM, HSCs occupy a relatively hypoxic niche compared to more differentiated lineages, particularly the MKs which shed mature platelets into the blood vasculature. This migration from hypoxic to normoxic environment increases the level of reactive oxygen species (ROS) in these cells, which has been indicated as a crucial signaling molecules (34). In fact increased oxygen level and augmented expression of ROS generating enzymes show positive correlation with megakaryocyte development, corroborating ROS to serve a critical promoter of megakaryopoiesis (35-37). ROS has been suggested to function at multiple levels which include increasing the level of tyrosine-phosphorylation on some proteins or by inhibition certain tyrosine phosphatases or by maintenance of ERK activation or modulation of cyclin levels or increasing expression of crucial TFs or promoting differentiation associated apoptosis (35, 38-40).

K562 cells represent a bi-potential cell line which can be differentiated to either MK-like cells or Erythroblast-like cells depending on differential pharmacological stimulation (41). Supplementation of Phorbol esters like Phorbol-12 myristate-13 (PMA) in growth media of K562 cells drives differentiation towards megakaryocytes, inducing cellular and molecular changes akin to MKs *in vivo*, through activation of identical signaling axes (42-50). In addition to this K562 is permissible to DENV replication and is therefore a good model system to study the effect of DENV infection on Megakaryopoiesis (51). In this report we show results suggesting that when DENV infected MEPs differentiate into megakaryocytes the intracellular viral replication is promoted, by a mechanism that is not clear yet. This enhancement of viral replication affects key signaling pathways that are known to be crucial for maturation of megakaryocytes, without any significant effect on their survival. DENV infection is known to induce an accumulation of ROS in the host cells. K562 cells induced to differentiate into megakaryocytes by treatment with PMA also accumulate ROS. However surprisingly, infected cells undergoing differentiation accumulate significantly less ROS when compared to uninfected controls, suggesting an active suppression of ROS by viral replication. The implications of these observations with respect to Megakaryopoiesis and DENV replication have been discussed.

## Results

### K562 cells recapitulate major events in Megakaryopoiesis upon PMA-treatment

Differentiation of Mk from an MEP follows elaborate cellular changes involving a dramatic increase in cell size and endomitosis with consequential formation of a multi-lobed, polyploid nucleus as depicted in the schematic figure (Fig. 1A.). K562 cells mimic a bi-potential Megakaryocyte-Erythrocyte progenitor (MEP) and depending upon pharmacological treatment can be differentiated into cells of either Mk-lineage (Phorbol esters) or Erythroblast-lineage (Sodium Butyrate or NaB) (41). Upon supplementation of Phorbol-12 Myristate-13 acetate (PMA) in culture media, K562 cells stopped proliferation and enlarged in size (data not shown). Further, the cells underwent endomitosis leading to generation of polyploid cells harboring multilobed nuclei (Fig. 1B.). When analyzed by flow cytometry, the side scatter of PMA-treated K562 cells showed a significant increase indicating accumulation of cytoplasmic granules (data not shown). The platelet plasma membrane is derived from that of the Mk mother cell and during differentiation platelet-specific surface markers are expressed on the Mk surface. As a corroboration of megakaryopoiesis induced by PMA-treatment, the surface expression of three platelet specific markers were quantified in K562 cells and the result showed a dramatic increase in the level of CD41/61, CD61 alone and a significant increase in number of cells positive for CD42b expression (Fig. 1C.). Analysis of the cell ploidy following PMA-treatment showed an almost complete disappearance of cells in the S-phase and emergence of polyploid cells with ploidy of 8N and 16N (Fig. 1D.). As a control for these changes being specific to K562, PMA-supplementation did not stall the growth or induce expression of platelet-specific surface markers in cells of the hepatoma cell line Huh7 (data not shown). Since treatment with NaB induces differentiation towards an Erythroblast-lineage, the differential expression of specific genes were compared between K562 cells treated with either PMA or NaB. The result showed a distinct pattern of gene expression induced by these pharmacological treatments differentiating the two lineages, especially with respect to transcripts of the gene GATA2, EBP42, GYPA and CD61 (Fig. 1E.).

**Figure 1:**
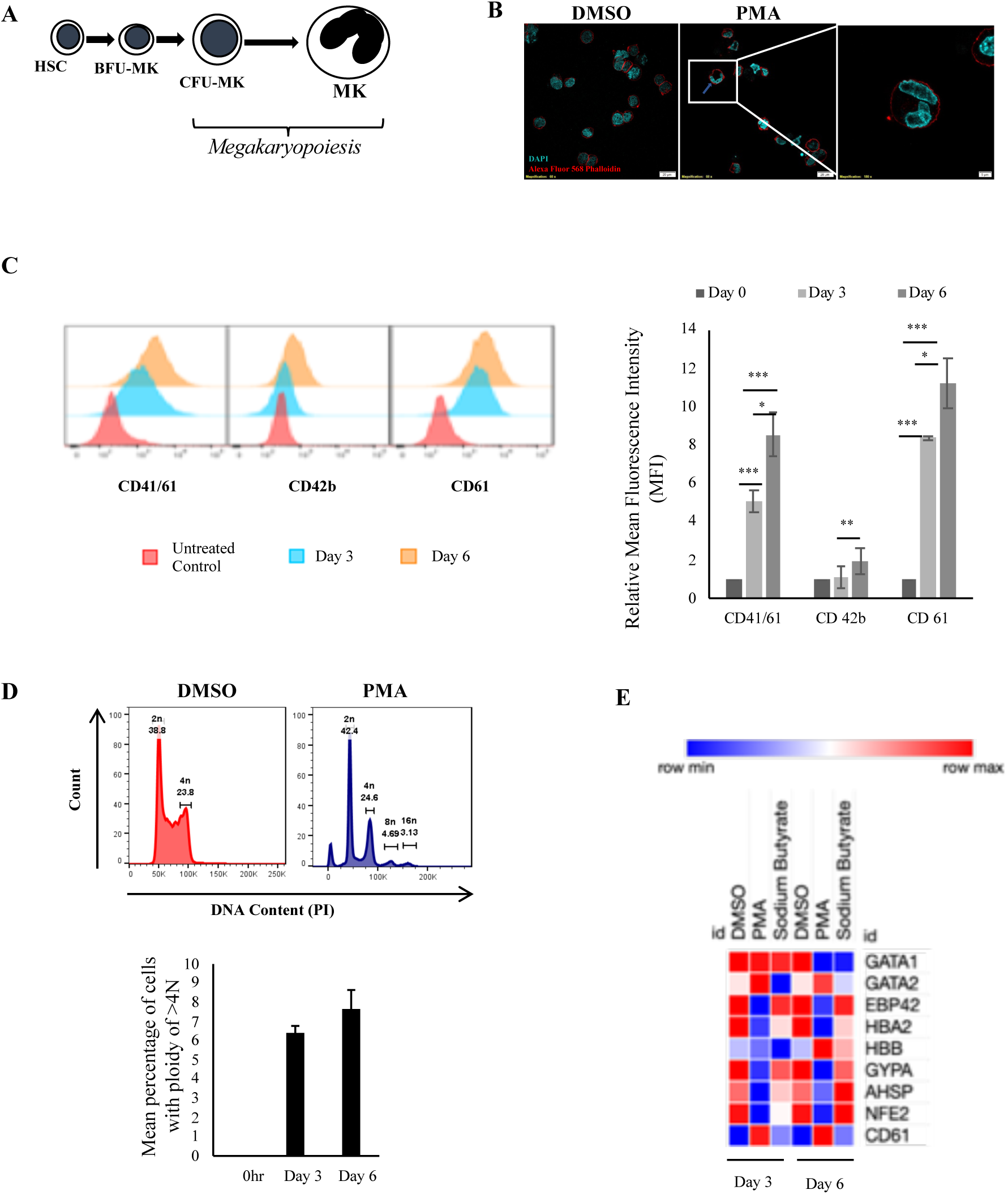
**Panel A: Schematic diagram of Megakaryopoiesis**: The schematic diagram shows some of the lineages that are produced during differentiation of Hematopoietic stem cells (HSC) to Megakaryocytes (Mk). The formation of multilobed nucleus has also been depicted. **Panel B: Visualization of endomitosis and multi-lobed nuclear formation following PMA-treatment**. K562 cells treated with either DMSO or 50 nM PMA for 6 days were stained with Alexa Fluor-568 Phalloidin, mounted onto glass slides using mountant supplemented with DAPI and visualized in a confocal microscope. The red color indicates the plasma membrane and blue color shows nuclei. The inset shows a cell with multilobed nucleus. **Panel C**: **Expression of platelet specific surface markers:** K562 cells treated with PMA for 0, 3 or 6 days were immune-stained for expression of the indicated surface marker protein using fluor-conjugated primary antibodies. The mean fluorescence intensity (MFI) corresponding to each day was quantified. For each surface marker the MFI at day 0 was arbitrarily set to 1 and those at day 3 and day 6, expressed as the relative fold change with respect to that. The error bars represent standard deviation. The significance was calculated by Student’s t-test (*, **, *** respectively indicate P-values <0.5, <0.1 and <0.01). **Panel D**: K562 cells treated with PMA for 0 or 3 or 6 days were fixed, permeabilized and stained for intracellular DNA using Propidium iodide (PI). The PI stain was quantified by flow cytometry and mean number of cells having ploidy of >4N was plotted. The error bars represent standard deviation. **Panel E:** The total RNA from K562 cells treated with either DMSO or 50 nM PMA or 1mM Sodium Butyrate (NaB) for 3 or 6 days was extracted and purified. The total RNA was reverse transcribed using random hexamers and the cDNA obtained used for real-time PCR estimation of mRNA transcripts from indicated genes. The C_T_ value corresponding to each was normalized to that of GAPDH. The normalized value at day 0 was taken arbitrarily as 1 and those at day 3 and 6 expressed as fold-change with respect to that.

### Differentiation of K562-MKs promote replication of Dengue virus

Differentiation into megakaryocytes is associated with arrest in cell cycle and other molecular changes. In order to analyze how these changes might affect the ongoing intracellular replication process, DENV replication in infected K562 cells that have been treated with PMA soon after infection, was compared to simultaneously infected cells treated with vehicle (DMSO). For this purpose, the level of DENV envelope protein accumulated was compared between these conditions at 3 or 6 days post-infection by immunofluorescence followed by analysis in a flow cytometer. The result showed that although the viral protein accumulated at 3 days p.i was comparable between PMA-treated and DMSO-treated cells, a dramatic difference between them was observed at day 6 (Fig. 2A.). This difference was observed using different MOI of infection, although due to saturation of immunostaining the difference was less dramatic at higher MOI (data not shown). In order to check if higher accumulation of viral protein correlates positively with viral replication, the level of intracellular viral RNA and infectious virus in culture supernatant was compared between infected K562 cells treated with either PMA or DMSO for 3 or 6 days. The results showed a similar higher accumulation of intracellular viral genomic RNA and extracellular infectious virus, between these two conditions at 6 days post-infection (Fig. 2B and 2C). During virus maturation the prM protein on assembled virion particles undergoes a cleavage by the host protease Furin, a step necessary for conferring infectivity to the secreted virus (52). Therefore the comparatively higher infectious titer in culture supernatant of PMA-treated cells can result from either higher secretion of virus (more virion particles without any change in their infectivity) or more efficient Furin cleavage brought about by the cellular and biochemical changes accompanying differentiation (same number of virion particles as secreted by DMSO-treated cells but with higher infectivity). To dissect these possibilities, the viral genomic RNA from culture supernatant of cells kept with either PMA or DMSO was extracted and purified and their relative level compared by real-time PCR using DENV specific oligonucleotide primers. The comparison showed comparatively more viral genomic RNA in the supernatant of PMA-treated cells, suggesting that the differentiating cells secrete more virus that are produced by higher replication efficiency in these cells (data not shown). However this indicated towards possibilities wherein the entry within differentiating cells might be facilitated e.g. through overexpression of viral receptor. In order to address this, uninfected K562 cells treated with either DMSO or PMA for 3 days were incubated with DENV inoculum and the viral entry compared by comparison of the internalized genomic RNA immediately after entry. The result showed a comparable entry into cells irrespective of no differentiation or differentiation for 3 days, implying that the promotion of replication occurs post-entry (Fig. 2D). As a negative control DENV-infected Huh7 cells did not exhibit any differential replication depending on treatment with PMA and DMSO (data not shown). Also, treatment of infected K562 cells with Na-butyrate which would initiate erythropoiesis, did not exhibit a similar promotion of the viral replication (data not shown).

**Figure 2:**
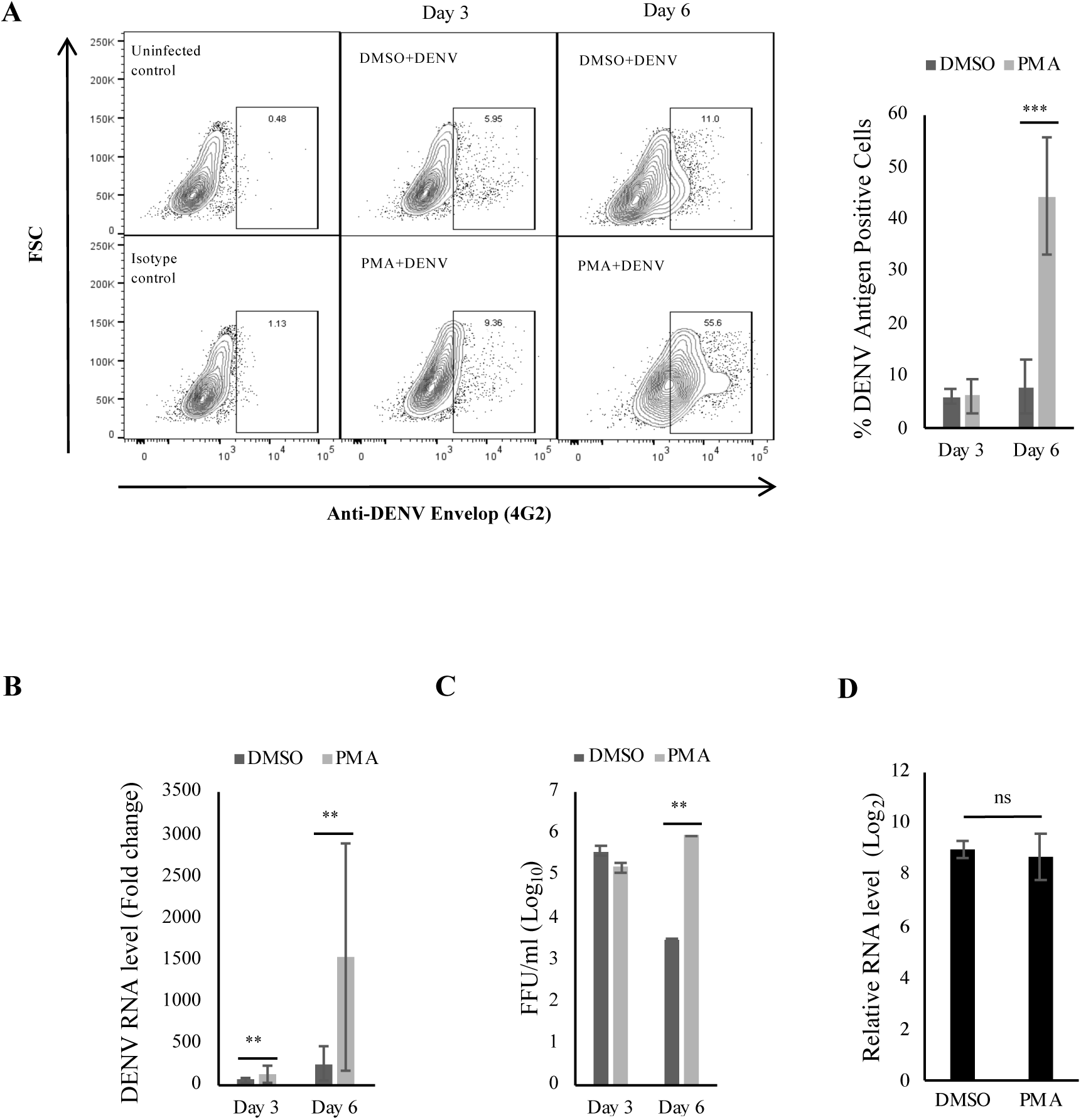
DENV replication is increased upon PMA treatment of infected K562 cells: **Panel A:** K562 cells infected with DENV at an MOI of 0.1 were maintained in growth media supplemented with either DMSO or 50nM PMA and fixed with PFA at 0 hour or 3 days or 6 days post-infection. The cells were permeabilized and immune-stained for intracellular DENV antigen and the fluorescence quantified by flow cytometry. The fluorescence in cells fixed at 0 hour was used to gate for DENV antigen positive cells. The number of antigen positive cells were quantified and plotted. The error bars represent standard deviation and the significance was calculated by Student’s t-test (*** indicates P <0.001). **Panel B**: Total RNA from K562 cells treated similarly as in panel A was extracted, purified and reverse-transcribed using random hexamers. The cDNA was used for real-time PCR comparison of DENV RNA level at 3 or 6 days to that at 0 hour, normalized to that of GAPDH mRNA. The value at 0 hour was taken arbitrarily as 1 and that at other time points expressed as fold-change over that. The error bars represent standard deviation and the significance was calculated by Student’s t-test (** indicates P <0.01). **Panel C**: The culture supernatant of cell infected with MOI of 0.1 and subsequently treated as in panel A was collected and used for quantification of focus-forming unit/ml (FFU/ml) on Vero cell monolayers. The FFU/ml for each sample was transformed to their logarithmic value to the base of 10 and plotted. The error bars represent standard deviation and the significance was calculated by Student’s t-test (** indicates P <0.01). **Panel D**: Uninfected K562 cells treated with either DMSO or PMA for 3 days were washed with PBS and incubated in either mock- or DENV inoculum (10 MOI) on ice for 2 hours. The cells were then transferred to 37°C and further incubated for 2 hours. Subsequently cells were washed, treated with 0.25% trypsin for 5 minutes and the total RNA extracted and purified. 1.0 μg of total RNA was reverse-transcribed with random hexamers and the cDNA used in real-time PCR for comparison of DENV genomic RNA normalized to GAPDH transcripts. The level in mock-infected cells was taken as 1 and that in DENV infected cells calculated as fold-change over that. The relative enrichment of DENV RNA compared to mock-infected cells was quantified, transformed to their logarithmic value to the base of 2 and plotted. The error bars show standard deviation and the statistical significance was calculated by Student’s t-test (ns= not significant).

### Cellular and biochemical changes imposed by DENV infection in differentiating K562 cells

Differentiation of Mk from HSCs is influenced by multiple cytokines, the most characterized among which is Thrombopoietin (TPO) which is secreted from liver cells and binds to the surface receptor c-Mpl. Interaction of TPO with c-MPL activates Janus kinase 2 (JAK2) tyrosine kinase initiating a cascade of phosphorylation events (Fig. 3A). Although an important cytokine, TPO is not indispensable for megakaryopoiesis since a knock-out of the gene encoding TPO (TPO^-/-^) is not lethal in mice, albeit with the platelet level being at 10% of that in wild-type animals. Phorbol esters like PMA are known to activate kinases like Protein kinase-C (PKC) that in its turn transduces the signal through nodes many of which are shared with TPO signaling. In response to viral infection mammalian cells secrete cytokines like Interferons that bind to cell surface receptors either in an autocrine or paracrine manner and induce an ‘anti-viral’ state within the cell. JAK2 is also central to signaling via the interferon receptor consequent to binding of type-I IFNs to the receptor (IFNR). This implied that megakaryopoiesis even when it is PMA-induced would potentially create an antiviral state within the cell and in that context it would be interesting to observe how DENV is still able to replicate to higher extent compared to undifferentiated cells. Since both of these pathways would also be potentially perturbed by a virus infection, we investigated the status of different members by immunoblotting, in cells which were stimulated to undergo PMA-dependent differentiation immediately after infection.

**Figure 3:**
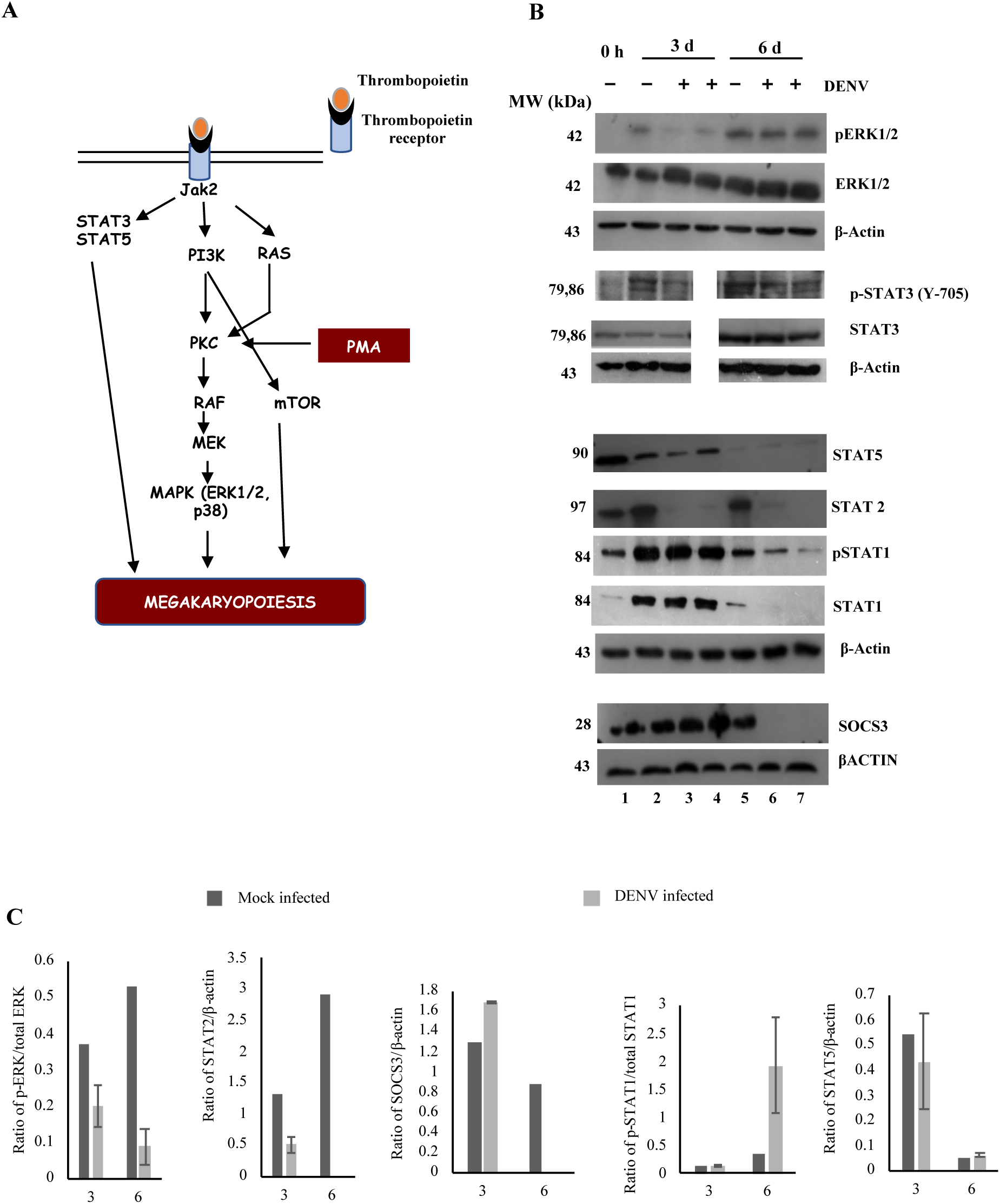
DENV replication in differentiating K562 cells interfere with PMA-induced signaling: **Panel A:** Schematic showing comparison of signaling activated upon TPO binding to c-Mpl and by PMA. **Panel B: Immunoblot for signaling protein activated by PMA in K562 cells**. K562 cells either mock-infected or DENV-infected were incubated in PMA-supplemented growth media and the total protein extracted after 3 or 6 days. The protein in each lysate was denatured, resolved in SDS-PAGE and immunoblotted for proteins indicated at right. Immunoblot for b-actin was performed for each batch of lysate used. The relative molecular mass of the marker protein is indicated on the left. **Panel C: Quantification of immunoblot bands in panel B**. the intensity of the immunoblot band for indicated proteins were quantified, normalized to that of b-actin and the ratio calculated before plotting. The error bars represent standard deviation of the ratio derived from at least two independent experiments, performed in multiple replicates.

Phosphorylation of the MAPK ERK1/2 plays a crucial role in MK development and expectedly we observed an increase in the ratio of p-ERK/ERK both subsequent time points when compared to day 0 (Fig. 3B, compare lanes 2 and 5 with lane 1). Interestingly, we observed that DENV infection to reverse the PMA-induced ERK1/2 phosphorylation at 3 days post-infection (Fig. 3B, compare lanes 3 and 4 with lane 2). However, by day 6 the phosphorylation of ERK1/2 recovered significantly in infected cells when compared to uninfected controls (Fig. 3B, compare lanes 6, 7 with lanes 3, 4). Nonetheless, the ratio of p-ERK/ERK at 6 days post-infection was still lower in virus infected cells when compared to uninfected cells treated with PMA for a similar duration (Fig. 3B, compare lanes 6 and 7 with lane 5). In a manner opposite to that of ERK1/2, a dephosphorylation of p38 MAPK has been similarly reported to be crucial for MK development. We did not observe any modulation in the phosphorylation status of p38 in DENV-infected cells treated with PMA for 3 or 6 days (data not shown).

In addition to MAPKs we compared the status of different signaling protein in the JAK-STAT pathway in uninfected or DENV-infected K562 cells treated with PMA. While some effect of virus infection was observed with respect to a few of these signaling mediators, most of them were unaffected. A few more JAK-STAT pathway intermediaries were probed by immunoblotting but remained undetectable under presently used conditions (SOCS1, PIAS1, PIAS3 and PIAS4). When compared to 0 hour the ratio of p-STAT1/STAT1 showed a significant decrease upon PMA treatment at both time points (Fig. 3B, compare lanes 2 and 5 with lane 1). However, although DENV replication in PMA-treated cells did not alter this ratio at 3 days p.i., by 6 days p.i. the ratio showed a dramatic increase in infected cells when compared to uninfected controls (Fig. 3B, compare lanes 3, 4 with 2 and lanes 6, 7 with lane 5). Further, at 6 days p.i. the total STAT1 protein level was also reduced to undetectable level in infected cells (Fig. 3B, compare lanes 6 and 7 with lane 5).

STAT3 has been suggested to promote Megakaryopoiesis and in support of this we observed an increase in ratio of p-STAT3/STAT3 at 3 days after PMA treatment, which was maintained till 6 days (Fig. 3B, compare lanes 2 and 5 with lane 1). However in DENV infected cells treated with PMA, the ratio of p-STAT3/STAT3 at both 3 and 6 days post-treatment was significantly lower when compared to uninfected controls (Fig. 3B, compare lane 3 with lane 2 and lanes 6,7 with lane 5). The lane 4 in this immunoblot is missing due to the technical reason of sample insufficiency.

Although STAT2 protein level is not known to be modulated by PMA-treatment, it is known to be degraded in host cells following DENV infection, in a NS5 dependent manner (Ref). We did not observe any alteration in STAT2 levels with PMA-treatment of uninfected cells at either time point (Fig. 3B, compare lanes 2 and 5 with lane 1). As expected however, the protein was reduced to undetectable level in DENV infected cells. (Fig. 3B, compare lanes 3,4 with lane 2 and 6,7 with lane 5).

A downregulation of STAT5 protein has been shown to be essential for promoting development of MKs. In concurrence with that, we observed a reduction in the level of this protein upon PMA-treatment starting from day 0 to almost undetectable level by day 6 (Fig. 3B, compare lanes 2 and 5 with lane 1). No additional modulatory effect was however observed by DENV infection of PMA-treated cells (Fig. 3B, compare lanes 3,4 with lane 2 and 6,7 with lane 5).

An activation of the JAK-STAT signaling upregulates inflammation, which eventually leads to expression of a family of genes that suppresses this pathways and are called as Suppressor of cytokine signaling or SOCS1, SOCS2 and SOCS3. In accordance with a crucial role for activated JAK-STAT pathway in Mk development, expression of the SOCS3 protein has been shown to have a negative effect on megakaryopoiesis. However, the SOCS3 level following PMA-treatment was observed to be unchanged at either time points (Fig. 3B, compare lanes 2 and 5 with lane 1). Surprisingly, in DENV-infected cells SOCSs was observed to be reduced to undetectable level by day 6 (Fig. 3B, compare lanes 6 and 7 with lane 5).

These results implied that DENV infection might be selectively affecting the innate antiviral and inflammatory pathways in differentiating K562 cell through degradation of STAT2 and SOCS3 level and interference with STAT3 phosphorylation.

### DENV infection inhibits polyploidy and accumulation of ROS in differentiating cells without affecting apoptosis

The biochemical changes associated with Megakaryopoiesis affect the cellular modifications. Therefore we quantified the PMA-induced surface marker expression and polyploidy was compared between uninfected or DENV-infected cells treated with PMA. As above, cells that were either mock-infected or infected with DENV were treated with PMA-supplemented growth media soon after infection and analyzed at 3 or 6 days post-treatment. The results showed a significant reduction in the surface expression of CD41/61, CD42b and CD61 in DENV infected cells (Fig. 4A). Phosphorylation of the ERK1/2 MAPK has been shown to be an critical modulator of polyploidy during Megakaryopoiesis. Since we observed a suppression of ERK1/2 phosphorylation at the early time point post-infection, a comparison of the percentage of polyploid cells at both time points was quantified by PI staining. As expected the result showed a significant reduction in the percentage of polyploid cells generated following DENV infection (Fig. 4B). Generation of platelet from megakaryocytes has been demonstrated to be dependent on induction of apoptosis in the mother cell and proceeds via a unique method of intrinsic apoptosis. In accordance with PMA-induced Megakaryopoiesis in K562 cells we observed a significant increase in Annexin-V positive cells upon PMA-treatment (data not shown). However, a comparison of mock-infected and DENV-infected cells treated with PMA for either 3 or 6 days did not show any difference in the proportion of Annexin-V positive cells (data not shown). However surprisingly, a comparison of Caspase3/7 activation showed significant increase in cleavage activity in virus infected cells when compared to uninfected controls at 6 days p.i. (Fig. 4C).

**Figure 4:**
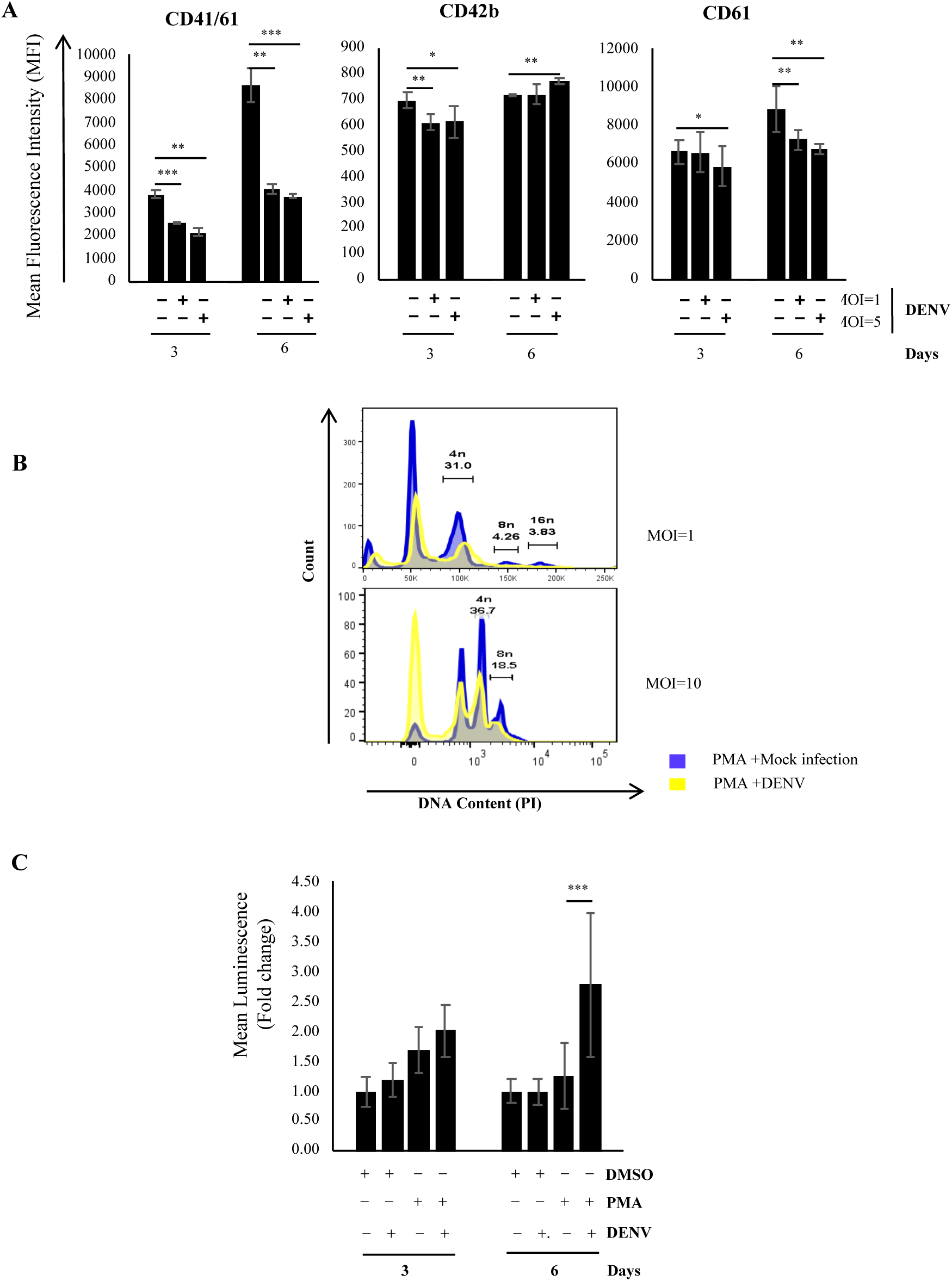
Effect of DENV replication on cell surface marker, polyploidy and apoptosis in differentiating K562 cells. **Panel A: Suppression of platelet-specific surface marker protein**. K562 infected with DENV at two different MOI, were incubated in PMA-supplemented growth media for 3 or 6 days. Subsequently the cells were immune-stained for the indicated surface marker proteins and the fluorescence quantified by flow cytometry. The mean fluorescence intensity was plotted. The error bars show standard deviation derived from at least two independent experiments in multiple replicates. **Panel B: DENV replication inhibits endomitosis**. DENV infected K562 were incubated in PMA-supplemented growth media for 3 or 6 days. Subsequently the cells were fixed and stained with propidium iodide and the fluorescence intensity measured by flow cytometry. The number of cells with ploidy of >4N were quantified and plotted. The error bars show standard deviation of the mean from two independent experiments performed in multiple replicates. **Panel C: Caspase cleavage assay in differentiated DENV infected cells**. Mock-infected or DENV-infected K562 were incubated in either DMSO or PMA-supplemented growth media for 3 or 6 days. Subsequently the cells were incubated with an substrate that can be cleaved by intracellular Caspase3/7 to generate a luminescent product. The luminescence was measured in a luminometer and plotted. The total luminescence at day 3 in DMSO-treated uninfected cells was arbitrarily taken as 1 and the rest plotted as fold-change over that. The error bars show standard deviation of the mean from two independent experiments performed in multiple replicates.

Intracellular ROS is known to be generated during megakaryopoiesis and an accumulation is essential for differentiation (47). DENV replication has been shown to induce generation of reactive oxygen species in infected cell (53). An estimation of the ROS accumulated in K562 cells as a result of DENV infection showed slight accumulation at 6 days post-infection (Fig. 5A, compare columns 2 to 1 and 6 to 5). In a similar manner, when compared to levels at day 0, uninfected cells treated with PMA showed a significant accumulation of ROS at both 3 and 6 days post-treatment (Fig. 5A, compare columns 3 to 1 and 7 to 5). We expected the ROS level in DENV infected cells that have been treated with PMA to be higher than the uninfected PMA-treated controls. However, surprisingly DENV infected cells treated with PMA did not exhibit additional ROS accumulation at day 3 post-treatment (Fig. 5A, compare columns 3 to 4). Further, surprisingly at day 6 after PMA-treatment the ROS accumulation in DENV infected cells were remarkably lower compared to uninfected PMA-treated cells (Fig. 5A, compare columns 7 to 8). This suggested DENV replication in differentiating cells to impede accumulation of ROS. In corroboration we observed DENV infection to have an effect similar to that of N-acetyl cysteine a biochemical known to quench ROS levels (Fig. 5B). In conclusion, DENV infection interferes with surface marker expression in addition to significantly affecting accumulation of ROS and endomitosis which are associated with differentiation into megakaryocytes.

**Figure 5:**
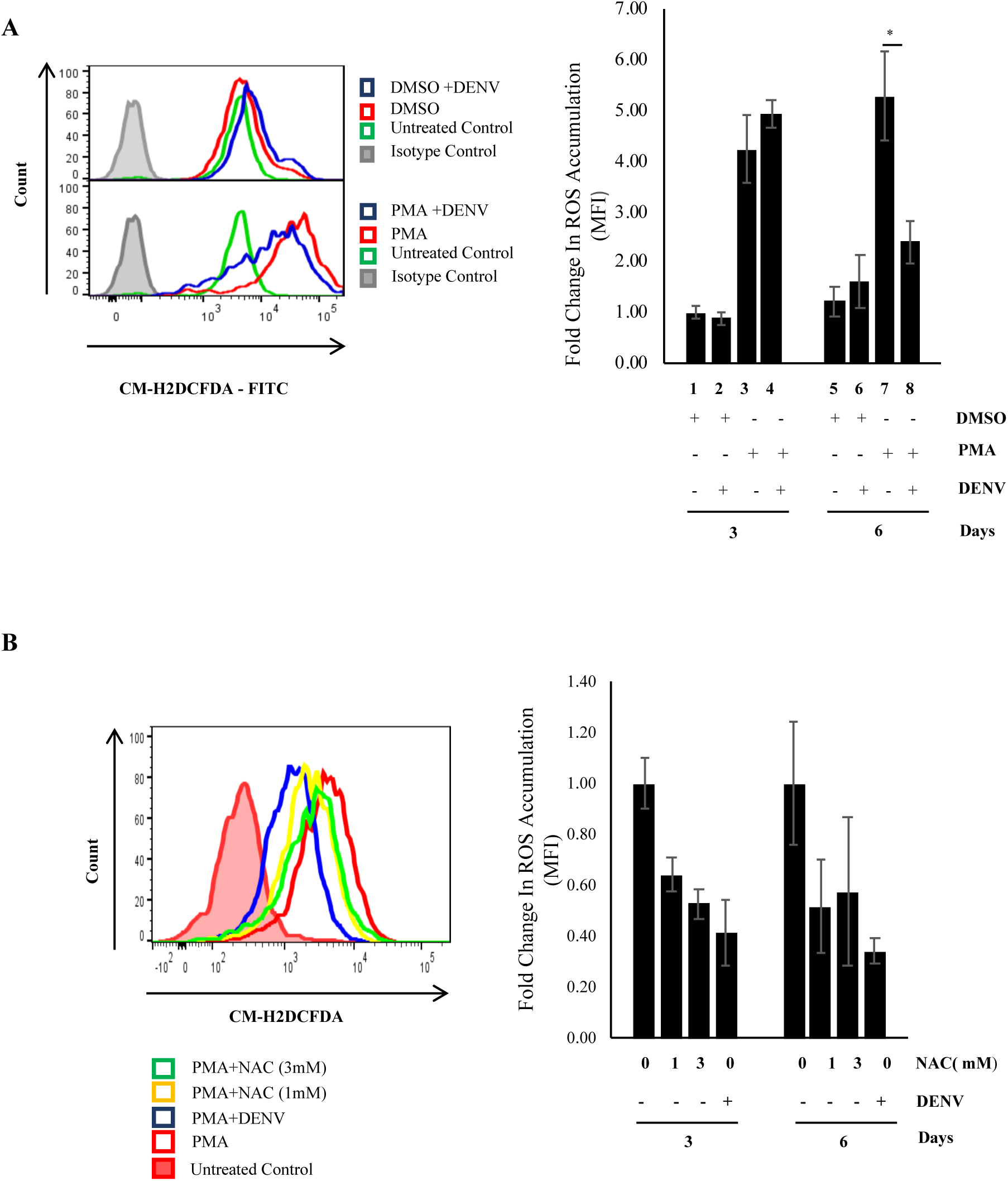
DENV infection suppresses ROS generated during PMA-induced Megakaryopoiesis in K562 cells: **Panel A**. Mock-infected or DENV-infected K562 cells were incubated in either DMSO or PMA-supplemented growth media for 3 or 6 days. Subsequently the cells were stained with H_2_DCFDA and the fluorescence quantified by flow cytometry. The mean fluorescence intensity in mock-infected and DMSO-treated cells at 3 days was arbitrarily taken as 1 and the rest calculated as fold-change over that. The error bars show standard deviation of the mean from two independent experiments performed in multiple replicates. **Panel B**. K562 cells either mock-infected or DENV-infected were immediately incubated in growth media supplemented with 50 nM PMA in addition to either 0 or 1 or 3 mM N-Acetyl Cysteine (NAC) as indicated. At 3 or 6 days post-infection the cells were stained with H_2_DCFDA and the fluorescence quantified by flow-cytometry. The mean fluorescence intensity in mock-infected cells treated with PMA and 0 mM NAC was arbitrarily taken as 1 and that in the rest expressed as fold change with respect to that. The error bars show standard deviation of the mean from two independent experiments performed in multiple replicates.

### PMA-induced transcriptome changes are reversed by DENV infection in differentiating cells

Megakaryopoiesis is associated with profound alterations in the transcriptome of differentiating cells. Similarly, innate antiviral response launched upon detection of a viral infection is also associated with changes in the pattern of expression of many genes. Therefore, in order to study the interaction between these two stimuli at the transcriptional level the transcriptome of mock-infected and DENV-infected cells was compared after 6 days of PMA-induced differentiation. For this purpose, K562 cells either mock-infected or DENV-infected were induced to differentiate using 50 nM PMA and the total RNA isolated for analysis of transcriptome by next-generation sequencing. The transcriptome of both uninfected and DENV-infected cells was compared to uninfected cells at 0 hour with respect to PMA-treatment. As shown in Fig. 6A, the transcriptome in uninfected cells underwent a profound change in more than 5000 genes showing different degree of deregulation. Interestingly, the comparison between uninfected and DENV-infected cells after 6 days of differentiation showed reversal in the direction of PMA-induced deregulation for a number of genes (Fig. 6B.). Since our earlier result showed that DENV infection suppressed ROS accumulation in differentiating cells, ROS-associated genes the PMA-induced deregulation trend of which was reversed by infection were analyzed further. A heatmap of these genes showed that although alteration in transcript level was moderate for most of them, a few genes were dramatically affected (Fig. 6C). This included GCH1, FOXO3, GLRX5, ETHE1, CTNS, EPAS1, GFOD1 and SNCA (Fig. 6C). Among these transcripts corresponding to FOXO3, CTNS and EPAS1 were higher in infected cells compared to uninfected ones, whereas the others showed a relatively lower level in infected cells. FOXO3 and EPAS1 are transcription factors among which the former is known to be pro-apoptotic (54). Among the downregulated genes the protein corresponding to GCH1, ETHE1 and SNCA are enriched platelets (55-57). SNCA is also known to function in an anti-apoptotic manner although the specific role of this gene in apoptosis associated with megakaryocyte differentiation is not clear yet (58).

**Figure 6:**
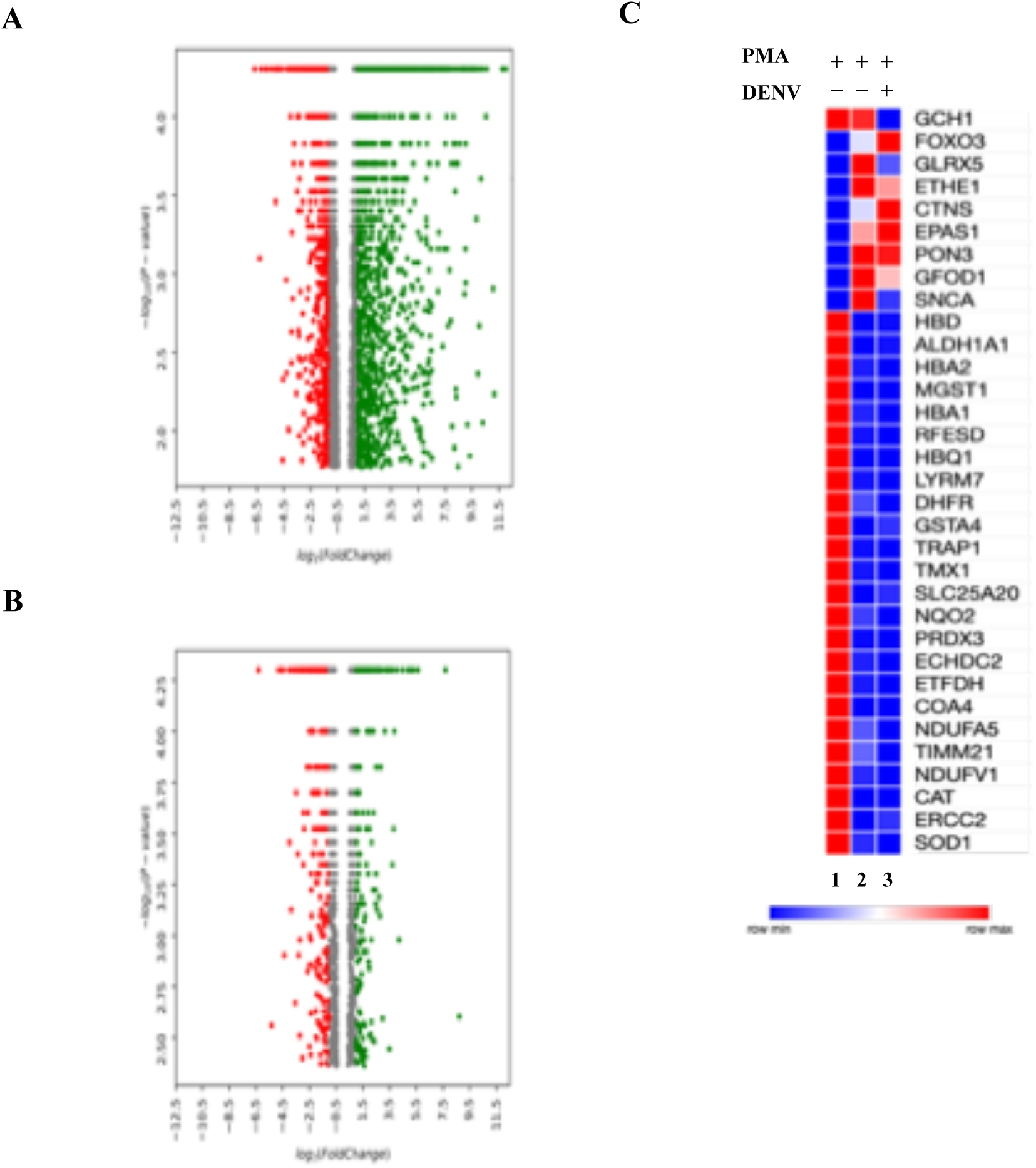
DENV replication interferes with PMA-induced transcriptome changes. **Panel A**: The transcriptome in K562 cells treated with 50 nM PMA for 6 days was compared to that after 0 hour by Next-generation sequencing. The genes differentially expressed after 6 days of treatment, either upregulated by ≥ 1.5 fold or downregulated by ≤ 0.5 fold with a P-value of ≤0.05, when compared to 0 hour were used to draw a Volcano plot prepared using in-house written Python script. The X-axis represents the logarithmic value to the base of 2 of the fold-change in gene expression. The Y-axis represents the negative logarithmic to the base of 10 of the P-value. **Panel B**: The transcriptome in K562 cells, either uninfected or DENV-infected and treated with 50 nM PMA or 6 days was compared to uninfected cells at 0 hour by NGS to find differentially expressed genes. The level of differential expression of each gene was then compared between uninfected and DENV-infected cells to draw the list of genes the PMA-induced deregulation of which is influenced by infection. A volcano plot was prepared from this list using in-house written Python script. The X-axis represents the logarithmic value to the base of 2 of the fold-change in gene expression. The Y-axis represents the negative logarithmic to the base of 10 of the P-value. **Panel C**: **Heatmap of genes associated with ROS-pathway**. The genes that are associated with the ROS-pathway in panel B were selected and enriched using the Metascape software. The PMA-induced fold-change in the transcript level of these genes after 6 days when compared to uninfected cells treated with PMA for 0 hour (column 1) in either uninfected cells (column 2) or DENV-infected cells (column 3), was represented by a heatmap prepared using the Morpheus algorithm (https://software.broadinstitute.org/morpheus). The colors represent the trend of deregulation corresponding to each gene shown on the right.

## Discussions

DENV replicates in multiple cell types and causes a range of pathological symptoms. Thrombocytopenia, a prominent DENV associated pathological symptom is a consequence of both impaired biogenesis and altered stability of platelets. Megakaryocytes or platelets mother cells in the bone marrow are known to be permissive for DENV replication, which affects platelet biogenesis. In this article, we have explored the cellular and molecular effect of DENV infection on Megakaryocyte development using the phorbol ester PMA induced-differentiation model in human K562 cells. Our results show that differentiation of infected K562 cells promote the replication of intracellular virus by an unknown mechanism and viral replication interferes with cell signaling pathways crucial for Megakaryopoiesis, particularly involving the MAPK ERK1/2 and different factors in the JAK-STAT pathway. However high level of viral replication does not affect the survival of the cells. PMA-induced differentiation of K562 cells, as with *in vivo* Megakaryopoiesis, is associated with accumulation of intracellular ROS and our results show that DENV replication suppresses the accumulation of this signaling molecule by an unknown mechanism. Replication by multiple viruses is known to increase ROS in the host cell we report an unique observation about DENV replication suppressing ROS accumulation under specific conditions in the host cell.

The absolute dependence of virus life cycle on host biochemical pathways implies a profound influence of host cell metabolic status on viral replication. Depending on the virus being studies, PMA a known activator of protein kinase C, is known to have either positive or negative effect on virus replication (59-62). On the other hand, viral infection of human monocytes can induce differentiation of these cells into Dendritic cells although the implication of this is not clear yet (63). Cell cycle arrest in PMA-treated K562 cells happens through upregulation of p21 and p27 in a protein kinase C (PKC)-dependent but p53-independent manner (64). Inducing an arrest in the host cell cycle is quite common among human viruses with beneficial effect on their replication (65). The entry of DENV is known to be influenced by the cell-cycle stage of the host cell, albeit in a cell-type dependent manner (66, 67). Here we observed increased intracellular replication in K562 cells that have stopped dividing in response to PMA, although the halt in cell division does not lower the metabolic activity as is obvious from increase in cell size and other cellular changes. In a similar manner Parvovirus B19 has been observed to infect cells of erythroblast origin with replication of the viral genome being promoted by differentiation of these cells into erythroblasts (68). Although it is not clear yet if the PMA-induced cell cycle stall is responsible for increasing intracellular virus replication, single-cell transcriptomics study by Zanini and coworkers suggested the cell cycle status of the host cell to have no effect on DENV genome replication (69). In addition to DENV, Mk cells are known to support replication of other viruses like the Human Immunodeficiency Virus and Hantavirus, with the former inducing apoptosis in addition to reducing the surface expression of c-Mpl (70-73). Interestingly, PMA-induced differentiation of Hantavirus infected megakaryocytes augmented virus replication, as observed in this study (72).

PMA induces rapid and profound changes in host gene expression and therefore it is possible that specific promoters of viral replication are overexpressed upon PMA treatment as observed for Hepatitis C virus which is benefitted by the liver cell enriched microRNA-122 (74). PMA-induced differentiation correlates with increase in granularity of K562 cells, probably pertaining to platelets granules that are synthesized in the megakaryocyte mother cell and transferred into platelets (75). Therefore the rate of protein synthesis in these cells can be expected to be enhanced for synthesis of the proteins that will form part of these granules. In face the protein content of K562 cells is significantly increased during PMA induced differentiation (76).It is possible that translation of viral proteins benefits from this overall increase in rate of protein synthesis. The DENV genomic RNA of DENV has a 5’ end type-I cap identical to host mRNAs that helps in recruiting ribosomes in addition to evading innate antiviral arsenal (77). A role for enhanced host protein synthesis benefiting viral translation is corroborated from our observation that the entry of DENV is not higher in differentiated cells. Another possibility is that differentiation leads to suppression of specific anti-viral genes, thereby benefiting the replication.

Although TPO is a cytokine crucial for platelet biogenesis and mice knock-out for either the cytokine or its receptor show drastically reduced levels of both mature MKs as well as circulating platelets, residual level of both MKs and platelets in these mice suggest other factors to also contribute in MK development and platelet formation (78, 79). Receptor interaction of TPO activates JAK2 kinase leading to activation of the MAPK ERK1/2, an important regulator of cell cycle and proliferation that plays a crucial role in Mk maturation (80, 81) Previous studies of Mk development using K562 cells showed PMA to directly activate protein kinase C (PKC) which in turn induces ERK1/2 phosphorylation, an important signaling event even for *in vivo* Megakaryopoiesis (26, 50, 82, 83). A concurrent inhibition of the MAPK p38 on the other hand supports MK development, suggesting this kinase to oppose or retard ERK1/2 mediated changes (22). Dengue infection can activate the p38 MAPK which is responsible for enhancement of pro-inflammatory cytokines and apoptosis (84, 85). We observed a reduction in the level of PMA-induced ERK1/2 phosphorylation in DENV-infected cells, probably causing inhibition of endomitosis and platelet-specific surface marker protein expression, although a concomitant increase in p38 phosphorylation was not observed. Previous reports have conclusively shown DENV NS5 protein mediated degradation of the host STAT2 infection as a major mechanism for suppression of the host innate-antiviral pathway (86). We observed additional regulation of a few other members of the JAK-STAT pathway, namely STAT1, STAT3 and STAT5, although it is yet clear if this is restricted to PMA-treated K562 cells or is a general effect occurring in other cells too. STAT1 regulated by the transcription factor (TF) GATA-1 plays a specific role in endomitosis by controlling expression of cell cycle genes like CCND1, CCND2 and CCNE2TFs through the TF RUNX1 (87). PMA-induced Mk development in K562 has not been reported earlier to mimic the *in vivo* differentiation with respect to regulation of STAT3 and STAT5. In a manner corroborative of *in vivo* development, we observed a gradual reduction of STAT5 level in parallel with progression of differentiation (24). STAT3 is phosphorylated initially on Tyr-705 which triggers dimerization (88). In addition to this STAT3 can have other post-translational modifications e.g. phosphorylation of S-727 or acetylation of K-685, although induction of these modifications by PMA in K562 cells is not clear yet (89, 90). In our study we observed DENV infection to reverse the PMA-induced STAT3 phosphorylation on Tyr-705, possibly through inhibition of the respective kinase or activation specific phosphatases. In addition to phosphorylation STAT3 function can be also suppressed by Ubiquitin mediated degradation or interaction with proteins that are activated downstream of the JAK-STAT pathway in a negative feedback loop e.g. protein encoded by PIAS and SOCS genes (91-93). In fact SOCS3 is known as a specific inhibitor of STAT3 signaling pathway, for which we expected an upregulation in infected cells (90). Unexpectedly however, we observed a reduction in SOCS3 protein level by DENV infection for which either mechanism or implication with respect to viral life cycle is still unclear. One of the potential role for STAT3 in Megakaryopoiesis is driving accumulation of intracellular ROS levels through upregulation of NOX2 expression (94). Therefore, suppression of STAT3 phosphorylation might be directly affecting accumulation of ROS as observed here. Further characterization of this observation would probably shed more light on the implication of this regulation.

Reactive oxygen species or ROS, in the form of Hydroxyl radical (.OH) or Singlet oxygen (^1^O_2_) or Superoxide anion radical (O_2_. −), are generated either in response to stimuli including metabolic inflammation, exposure to pathogen or as secondary messengers in signal transduction pathway. The most well characterized sources of intracellular ROS are two enzymes associated with the mitochondria namely **N**icotinamide **A**denine **D**inucleotide-**Q**uinone (NADH-Q) reductase (Complex I), Q-cytochrome c oxidoreductase (Complex III) and the plasma membrane associated homologs of NADPH-oxidase or NOX (95-97). Intracellular ROS plays crucial role in regulation of protein tyrosine kinases and phosphatases through post-translational modification (98, 99). It can also activate multiple cellular pathways including MAPK, NFkB, Cell cycle genes (100). Intracellular ROS can either promote apoptosis by inducing damage to the mitochondrial membrane or inhibit it through oxidation of catalytic site Cysteines on executioner proteases like Caspase-3 (101). In our study suppression of ROS level coincided with activation of Caspase-3 cleavage activity, although the mechanism would need further investigation to draw a direct correlation between them.

In the BM, pluripotent hematopoietic stem cells (HSCs) reside in a relatively hypoxic area while during differentiation megakaryocytes relocate to more oxygen-rich region near blood vessels from where platelets can be shed into the vascular circulation (102). ROS serves a critical role in megakaryocyte formation since a low level of these signaling molecules in differentiating HSCs disfavor generation of (**M**egakaryocyte-**E**rythrocyte **p**rogenitors) MEP and in favor of (**G**ranulocyte-**M**onocyte **p**rogenitors) GMP (103). Intracellular ROS levels are regulated through the interplay of generators (like NOXs) and mitigatory pathways (like the KEAP1-NRF2 pathway) (37, 104). PMA has been shown to induce accumulation of ROS in K562 cells and ROS has been shown to be responsible for polyploidy and a reduction of ROS by NAC directly impacts polyploidy (43, 47). Accumulation of ROS activates the KEAP1-NRF2 pathway and activation of many genes that can suppress the cellular ROS levels. ROS is produced by cellular responses like Unfolded-protein response (UPR) or innate-antiviral pathway (105, 106). We observed a relatively higher level of intracellular ROS with respect to uninfected cells in K562 cells that are not undergoing differentiation. Since PMA-induced differentiation of K562 cells also causes ROS to accumulate in cells, we expected an additive effect of DENV replication on ROS accumulation in these cells. However, surprisingly DENV replication quenched intracellular ROS levels through an unknown mechanism. Although evidence from previous reports would strongly indicate this suppression of ROS to be major contributor to inhibition in platelet biogenesis by infection of Mk mother cells by DENV, it is still not clear if the viral replication in these cells is benefited from this or not. Future studies would be directed to address the exact mechanism through which DENV infection suppresses ROS accumulation in these cells and the physiological relevance of this suppression for the viral life cycle.

## Acknowledgement

The authors would like to thank Dr. Manjula Kalia, Regional Centre for Biotechnology, NCR Biotech Science Cluster for careful reading of the manuscript. The Translational Health Science and Technology Institute is acknowledged for providing all support for equipment and other infrastructure. The Core Technology Research Initiative (CoTeRI) in the National Institute for Biomedical Genomics, India is acknowledged for providing Next-generation sequencing facility. JK is supported by fellowship from the University grants commission. YR is supported from fellowship in the extramural research grant (BT/PR22985/MED/29/1168/2016) to SB from the Department of Biotechnology, Govt of India. This work was supported by extramural research grant (EMR/2016/005796) to SB from the Science and Engineering Research Board, Department of Science and Technology, Govt. of India.

## Materials and methods

### Cell culture, Drugs and Virus

K562 cells were procured from the American Type Culture Collection (ATCC) and cultured at 37°C, 5% CO_2_ in Iscove’s modified Dulbecco’s media (IMDM) supplemented with Penicillin (100 U/ml), Streptomycin (0.1 mg/ml) and 10%-Fetal Bovine Serum (FBS). Vero and C6/36 cells were procured from the cell line repository of the National Centre for Cell Sciences (NCCS), India and cultured respectively in Minimum Essential Medium (MEM) and L15 cell culture medium supplemented with 10% FBS. C6/36 cells were maintained at conditions of 28°C and atmospheric CO_2_.

Dengue virus serotype 2 (strain NGC) was amplified in C6/36 and the culture supernatant used as inoculum. The infectious titer of the virus was determined by Focus-forming unit (FFU) assay in Vero cells. For all infections, the inoculum was diluted in the respective culture media supplemented with 2% FBS, and the cells were incubated with the inoculum for 2 hours at the respective culture conditions of temperature and CO_2_ concentration, with intermittent rocking.

Phorbol-12 Myristate-13 acetate (PMA, Sigma Aldrich) and N-Acetyl Cysteine were diluted in recommended vehicles and stored as single use aliquots at -20°C. Fresh dilutions of N-Acetyl-L-Cysteine (Sigma) were used for every experiment.

### Focus-forming unit (FFU) assay

Vero cell monolayers in 24-well plate were infected with 10-fold serial dilutions of virus inoculum. The inoculum was incubated with cells for 2 hours at 37°C and 5% CO_2_ with intermittent rocking. The inoculum was then discarded and complete MEM added to each followed by incubation for 48 hours at 37°C and 5% CO_2_. Subsequently the cells were washed with phosphate-buffered saline (PBS), fixed with 2% para-formaldehyde (PFA) and permeabilized with PBS supplemented with 0.1 % Triton-X-100 and 1% bovine serum albumin (BSA). The permeabilized cells were washed with PBS and sequentially incubated anti-DENV primary antibody (dilution 1:400 of Mab8705, Millipore) and anti-Mouse Alexa-488 conjugated secondary antibody (dilution 1:500; ThermoFisher Scientific) both diluted in PBS supplemented with 1% BSA. After incubation with primary antibody, the cells were washed twice with PBS before addition of secondary antibody dilutions. The fluorescent foci were visualized and manually counted using a fluorescence microscope and the titer calculated as FFU/ml.

### Flow Cytometry Analysis

All flow cytometry analysis were performed in a BD FACS Canto II flow cytometer (BD Biosciences) under standard conditions and all raw data analyzed using Flow-Jo software. For immune staining of intracellular DENV antigen, K562 cells were washed with ice-cold PBS before fixation with PBS supplemented with 2% PFA and permeabilization with the same buffer containing 0.1% Triton-X-100. The permeabilized cells were sequentially stained with dilution of 4G2 (purified IgG2a mAb produced in the lab from Hybridoma-HB112) antibody and anti-mouse Alexa-488 conjugated secondary antibody. For immune staining of surface markers, cells were washed with PBS and incubated with dilutions of fluor-conjugated primary antibodies, specific to either human CD61-Phycoerythrin (PE) (BioLegend) or CD41/61-(Allophycocyanin) APC (BioLegend) or human CD42b-(BD Horizon brilliant violet 421) Bv421 (BioLegend). Antibodies were diluted in staining buffer (PBS with 1%BSA and 0.02% Sodium Azide) and cells were incubated for 1h at 4°C. Subsequently the cells were washed with PBS and analyzed by flow cytometry.

Propidium iodide staining for polyploidy analysis: Cells were harvested by centrifugation, washed twice with ice-cold phosphate-buffered saline (PBS) and fixed with 70% ethanol at 4°C for 30 min. After a PBS wash cells were treated with 100µl PBS supplemented with RNase-A (200µg/ml) and incubated at 37°C for 30 min. Cells were then washed using PBS and the genomic DNA stained with Propidium iodide (50µg/ml).

Annexin-V and PI staining: Apoptotic detection assays were carried out by surface labeling with the Ca2+ dependent phosphatidylserine-binding protein Annexin-V. Cells were harvested by centrifugation, washed twice with cold PBS and then re-suspended in 1x binding buffer(Apoptosis detection Kit I # 556578, BD Pharmingen) at 1x 10^6^ cells/ml. A 100µl aliquot was stained by addition of 5µl FITC Annexin V and 5µl PI. The reagents were mixed by gentle vortex and incubated for 15 min at RT in the dark. 400µl of 1x binding buffer was added to each tube and the cells analyzed by flow cytometry within 1 hr.

### RNA extraction and Quantitative RT-PCR

Total RNA from 1×10^6^ K562 cells were extracted using Trizol (Takara) and purified with Qiagen RNAeasy mini kit (Qiagen), as per manufacturer’s instructions. Viral RNA from cell-free culture supernatant was isolated using QIAmp viral RNA mini kit (Qiagen) as per manufacturer’s instructions. 1.0 ug of total RNA was reverse-transcribed with ImProm-II reverse-transcriptase (Promega, USA) and random hexamers (Sigma) as per manufacturer’s protocol, and the cDNA diluted with nuclease-free water before use for real-time PCR. Real-time PCR was performed with 2x SYBR mix (Takara) in a QuantStudio-6 Flex Real-Time PCR System (Applied Biosystems) using the default run program.

### Confocal microscopy

K562 cells were washed with PBS, fixed and permeabilized as described earlier for immunostaining purposes. The cells were stained with 2.5µl of Alexa Fluor-568 Phalloidin (Invitrogen, Thermo fisher scientific) at RT for 20 min to stain for F-Actin followed by PBS wash. Cells were then mounted onto glass slides using ProLong Gold Antifade Mountant supplemented with DAPI (Invitrogen, Thermo fisher scientific). The fluorescence was observed and imaged in a FLUOVIEW FV3000 confocal microscope (Olympus).

### Intracellular Caspase-3/7 activity assay

The intracellular caspase-3/7 activity was measured using Caspase-Glo 3/7 Assay (Promega). For this, 1 x 10^4^ cells in 50 μl were mixed with an equal volume of Caspase-Glo3/7 reagent in wells of 96-well luminometer plate. The plate was incubated at room temperature for 3 hours and the luminescence measured using a Orion II microplate luminometer (Berthold). The luminescence from culture media without cells was used as negative control.

### Measurement of intracellular Reactive Oxygen species (ROS)

The intracellular ROS accumulated was measure using CM-H2DCFDA probe (Life Technologies) as per manufacturer’s instructions. Briefly, K562 cells were washed with PBS and then incubated in PBS supplemented with CM-H2DCFDA as per manufacturer’s instructions. The cells were then incubated at 37 °C, 5% CO_2_ for 30 minutes and washed twice with PBS. The fluorescence intensity was quantified in a BD FACS Canto II flow cytometer (BD Biosciences) under standard conditions and the raw data analyzed using Flow-Jo software.

### Electrophoresis and Immunoblotting

Cells were harvested by centrifugation at 4°C and washed with ice -cold PBS. Cells were suspended in lysis solution [250 mM NaCl, 20 mM Tris-HCl (pH 7.4),, 1% Triton X-100, 1 mM EDTA, supplemented with 0.1mM PMSF and protease inhibitor cocktail] and incubated on ice for 15 minutes. The lysate was clarified by centrifugation (13 000×g for 10 min at 4°C) and the supernatant collected. The total protein in the lysate was estimated using Bradford’s method. 40µg of whole-cell lysate was denatured, resolved in a SDS-10% gel and transferred to nitrocellulose blotting membrane. The membrane was blocked with either 5% skimmed milk or 5% bovine-serum albumin (BSA) and incubated with dilution of respective antibodies as per manufacturer’s instructions (Cell Signaling technologies). The membrane was washed, incubated with dilution of HRP-conjugated secondary antibody and the bands visualized by ECL chemiluminescence.

### Next-generation sequencing

The total RNA from K562 cells either uninfected or infected with DENV at an MOI of 10 was extracted and purified. Total RNA from similar number of uninfected cells, harvested for RNA extraction immediately after addition of PMA was used as 0 hour control. The total RNA was subjected to ribosomal RNA depletion before a sequencing library was prepared for 2×100 bp end-paired sequencing in Illumina HiSeq2500 platform, generating 50 million reads per sample.

